# Coordinate based meta-analysis: new clustering algorithm, and inclusion of region of interest studies

**DOI:** 10.1101/2020.04.05.026575

**Authors:** Christopher R Tench

## Abstract

There are many methods of conducting coordinate based meta-analysis (CBMA) of neuroimaging studies that have tested a common hypothesis. Results are always clusters indicating anatomical regions that are significantly related to the hypothesis. There are limitations such as most methods necessitating the use of conservative family wise error control scheme and the inability to analyse region of interest (ROI) studies, which can be overcome by cluster-wise, rather than voxel-wise, analysis. The false discovery rate error control scheme is a less conservative option suitable for cluster-wise analysis and has the advantage that an easily interpretable error rate is estimated. Furthermore, cluster-wise analysis makes it possible to analyse ROI studies, expanding the pool of data sources. Here a new clustering algorithm for coordinate based analyses is detailed, along with implementation details for ROI studies.

## Introduction

The aim of understanding the brain processes in healthy and disease groups is a task that often involves imaging. Functional magnetic resonance imaging (fMRI) is used to probe function and explore differences between groups. Localised volumetric changes can also be explored using structural imaging and voxel-based morphometry (VBM). These very commonly practiced analyses in fact use a collection of methods, with different groups employing different methodologies that also evolve with time. With variability in methodology, often small subject samples, and common lack of type 1 error control, typical fMRI and VBM studies can be tricky to interpret individually, particularly in the absence of detailed prior models. Therefore, in a complex system such as the human brain, building a complete model of the effect being investigated is not easy. However, when multiple groups have tested a common hypothesis, a meta-analysis (MA) might summarise the overall evidence sufficiently to generate a model that can subsequently be tested.

Meta-analysis of brain imaging is arguably most completely achieved using image based meta-analysis (IBMA) (Salimi-Khorshidi et al., 2009), but this does require availability of whole brain data from each study; currently this is not the most common scenario. A practical alternative is coordinate based meta-analysis (CBMA) (Turkeltaub et al., 2002; Wager et al., 2003), which uses only the reported summary results published with almost all fMRI and VBM studies. Summary result tables include coordinates of peak effect in a standard space, such as Talairach (Talairach and Tournoux, 1988) or MNI, along with peak statistical effect magnitudes in the form of Z scores or *t*-statistics. Multiple algorithms are available to meta-analyse these effects indicating those that are most likely generalisable. CBMA results are clusters of voxels indicating anatomical regions directly related to the common hypothesis being tested by the studies analysed.

Many CBMA methods use a null hypothesis that the peak coordinates are uniformly distributed through the grey matter to discover statistically significant clusters, where the coordinate density is significantly higher than expected under the null (Turkeltaub et al., 2002). Other CBMA algorithms use the reported Z scores to estimate a voxel-wise statistical image by spatial extrapolation (Albajes-Eizagirre et al., 2018; Costafreda, 2012). While the effect size in most voxels is unknown from the reported summary results, these can be considered censored by the statistical threshold and estimated by assuming a random effect model where all effects are drawn from the same Normal distribution.

There are several limitations to these methods. Firstly, they are voxel-wise analyses meaning a family wise error (FWE) method of controlling the error rate is the best option. By comparison, the alternative false discovery rate (FDR) (Benjamini and Hochberg, 1995) control method is more sensitive and it provides the more relevant estimate: the proportion of significant voxels that might be falsely positive. Unfortunately, voxel-wise FDR is problematic because the false positive voxels can form false clusters that are not distinct from the true positive clusters. Another limitation is that only whole brain studies can be meta-analysed (Müller et al., 2018). Studies investigating a-priori specified regions of interest (ROI) violate the null hypothesis that coordinates are spatially random, and they cannot be considered voxel-wise censored by a global threshold for statistical significance; rather there may not have been an a-priori region analysed at the voxel. Finally, the random effect model used by some algorithms (Albajes-Eizagirre et al., 2018; Costafreda, 2012) requires that all studies report effects from the same Normal distribution, including those that are censored. Consequently, any study observing a true null effect, which is therefore censored, violates the random effect model assumption so the effect is incorrectly estimated based on the highly significant uncensored effects.

To overcome these issues one option is to analyse cluster-wise rather than voxel-wise; where a cluster is now a cluster of studies reporting in a similar anatomical location. Coordinate based random effect size (CBRES) MA (Tench et al., 2017) performs a conventional cluster-wise random effect meta-analysis on the reported effect sizes (standardised Z scores) and incorporates censoring. Unlike other methods using the effect size and censoring, CBRES implements a null hypothesis based on random coordinates to circumvent the assumptions of the random effects model. Being a cluster-wise analysis means that CBRES can use the less conservative and more interpretable FDR based method of type 1 error control called false cluster discovery rate. Here an update to the CBRES algorithm is reported. Coordinate based meta-analysis of networks (CBMAN) (Tench et al., 2020) also uses Z scores and works cluster-wise rather than voxel-wise, but analyses the data as a cooccurrence network. As with CBRES, CBMAN uses FDR to control type 1 error, however CBMAN tests a null hypothesis that the effect sizes in spatially separated clusters are uncorrelated. The actual magnitude of the effect sizes is not the important feature, rather the relative magnitude in pairwise clusters must be correlated indicating clusters of both spatial and statistical effect magnitude consistency across independent studies. Consequently, ROI studies can be considered a special case in pairwise clusters that include reported ROI effects but ignored otherwise. This expands the potential pool of studies that might be analysed beyond other methods.

In order to analyse cluster-wise rather than voxel-wise a suitable algorithm is needed to partition the coordinates into clusters. In CBRES a DBSCAN (Density-based spatial clustering of applications with noise) type algorithm (Ester et al., 1996) was used as it is fast enough to allow the thousands of coordinate randomisations necessary to achieve numerical convergence; at least 4000. In CBMAN a different clustering algorithm was employed, which was based on empirical distributions of clusters in CBMA. This algorithm is relatively slow, and not suitable for use with CBRES. Consequently, CBRES and CBMAN may produce slightly different results. In this report a new clustering algorithm is introduced that is fast enough to use with CBRES, and has some advantages over the previous algorithm. In addition, the implementation of ROI studies into CBMAN is reported. The software *will* be available for free (https://www.nottingham.ac.uk/research/groups/clinicalneurology/neuroi.aspx); please contact author for availability.

## Methods and analysis

### Clustering

In the first version of CBMAN (Tench et al., 2020) clustering involved finding peaks in study density by identifying voxels where the density, defined by a kernel density estimator, is a local maxima. The nearest coordinates from each study are assigned to the peaks to form clusters. One issue with this approach is the assumption that nearest coordinates can be from any direction, which may not be valid when clusters are not spherical. Furthermore, coordinates may be assigned to clusters that are unfeasibly far from the cluster peak, which requires an adjustment based on detecting outliers. Here a new kernel-based method is detailed that overcomes these issues.

### A new kernel density estimate based clustering algorithm for CBMA

The new cluster forming algorithm is based on DENCLUE (DENsitybased CLUstEring) (Hinneburg and Keim, 1998). The algorithm requires the spatial density of reporting studies, which is estimated by a weighted sum of Gaussian kernels, with width *σ*, over coordinates

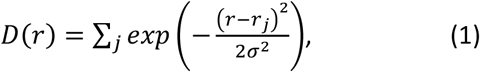

where *r_j_* indicates the coordinate from study *j* that is closest to position *r* where the density is estimated. An example of study density is depicted in figure (1) overlayed on a grey matter image. The kernel is determined by the empirical observation that typical clusters in CBMA have a spatial spread of coordinates with standard deviation between 3mm and 5mm with a large peak at around 4mm (Eickhoff et al., 2016b), which is modelled here with a truncated normal *N(4,0.4^2^)* distribution. The value of *σ* has been determined previously (Tench et al., 2020) such that a kernel with this width produces false density peaks rarely in simulated clusters. Here, *σ*=6.73mm is used, being the kernel width that produces 1% of false peaks in simulated clusters made up of 4 coordinates; 4 coordinates is the minimum number that defines a volume in three dimensions.

The smoothing kernel is designed to convert the reported coordinates into a smooth function consisting of isolated regions of high study density each with a single density peak (see figure (1)). Peaks are detected and coordinates assigned to clusters by using a recursive algorithm, which is initialised at each coordinate in turn and moves towards the density peak via neighbouring voxels in the direction of increasing density; see figure (2) and appendix for pseudo code. Multiple peaks are discovered, and multiple coordinates can lead to the same peak. It is also possible that multiple coordinates from the same study find the same peak, so only the closest to the peak is assigned to the cluster. Once the candidate clusters are identified, validity checks are performed: validity is defined by a cluster having a minimum number of independent studies contributing to it and having a minimum study density at its peak (as required by DENCLUE).

**Figure 1.**
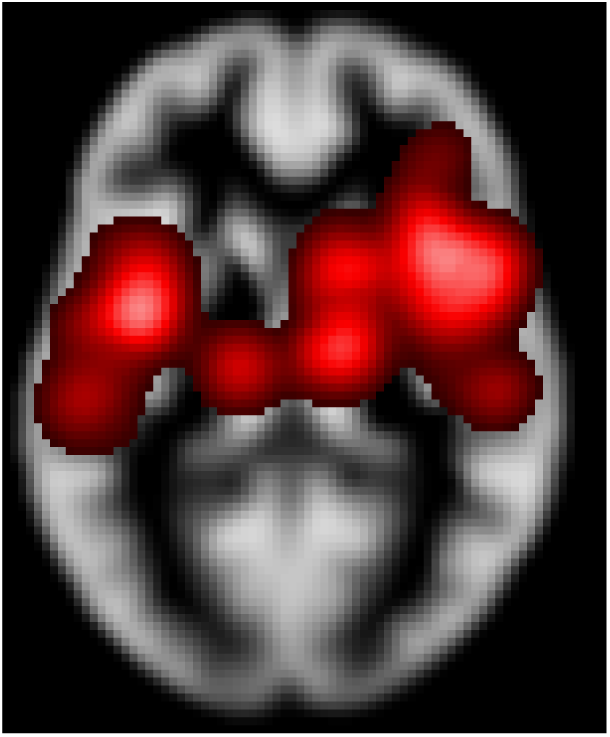
Example study density computed using equation (1).

**Figure 2.**
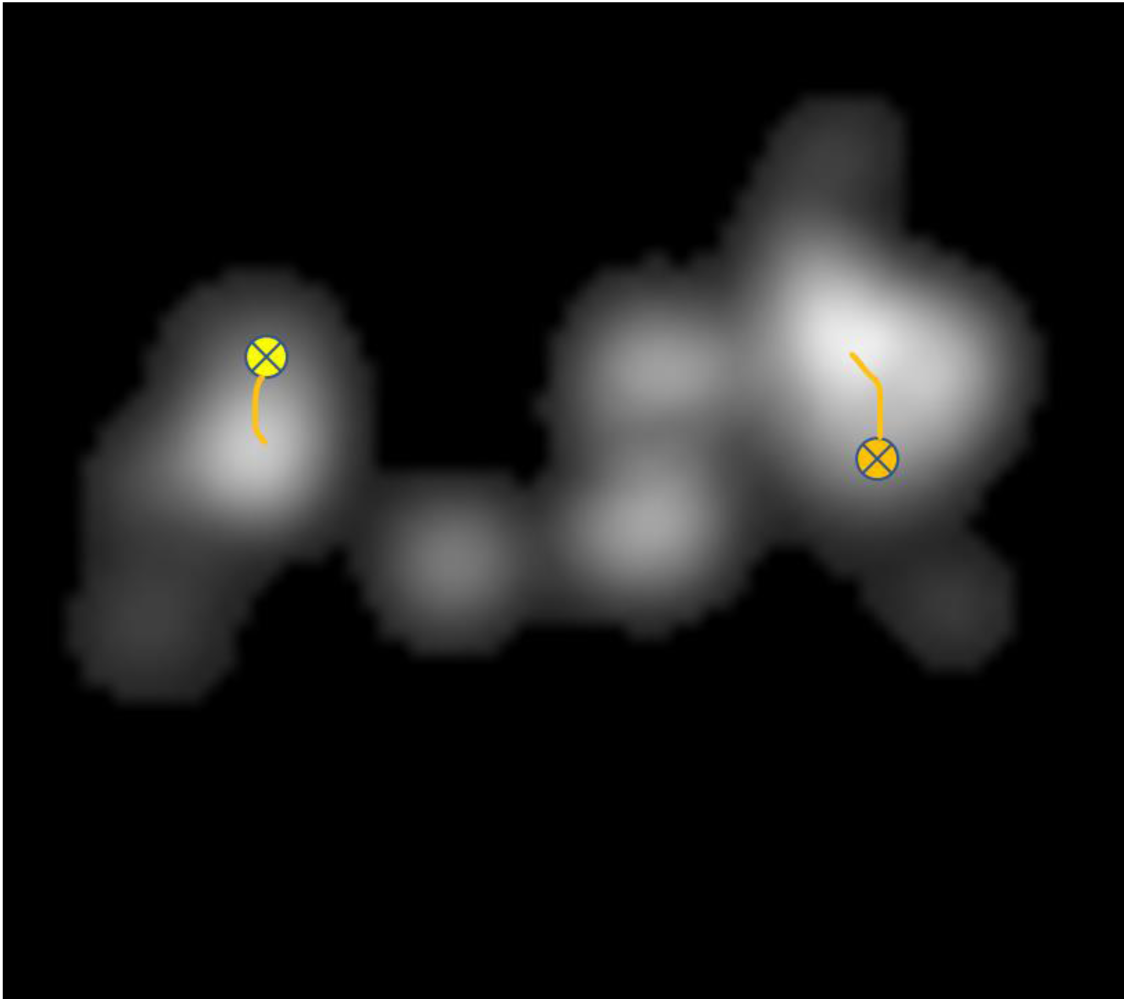
Depicting the study density with two coordinates (crossed circles) following the line of increasing density to the peak using the recursive algorithm given in the appendix.

**Figure 3.**
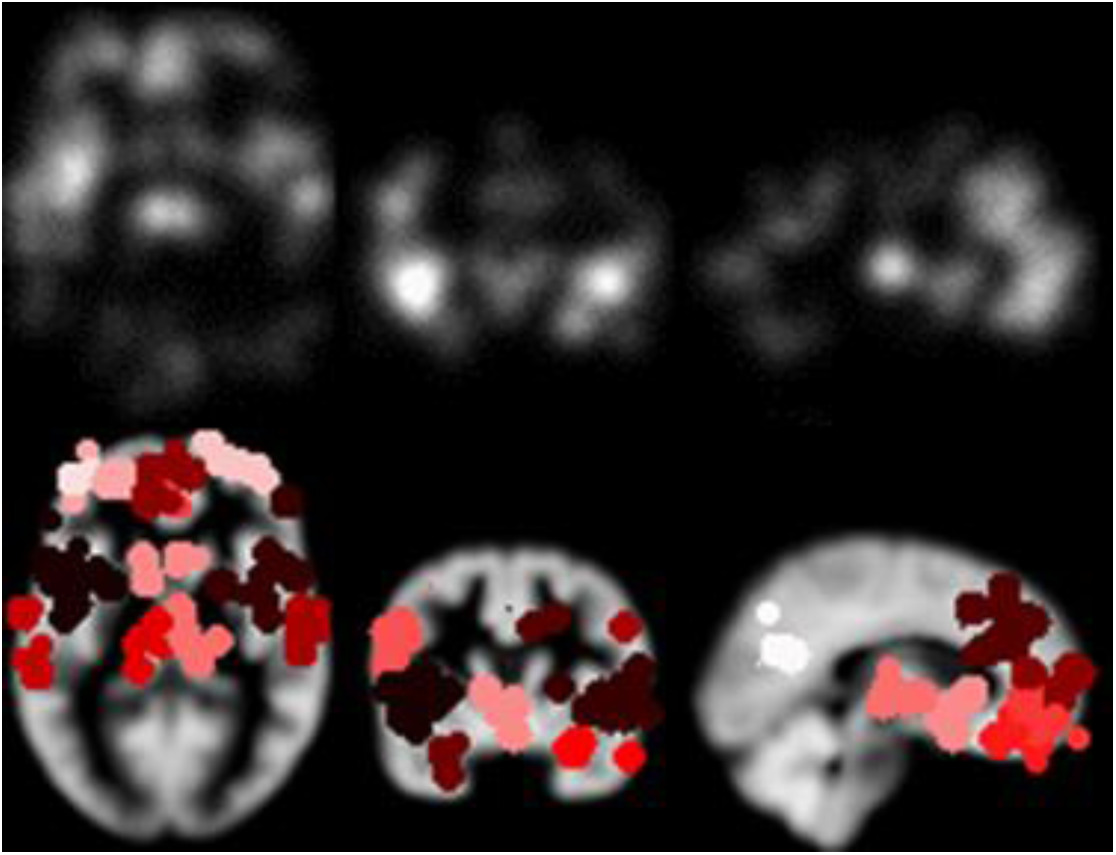
The study density estimated using equation (1) (top) and the resulting clusters (bottom).

**Figure 4.**
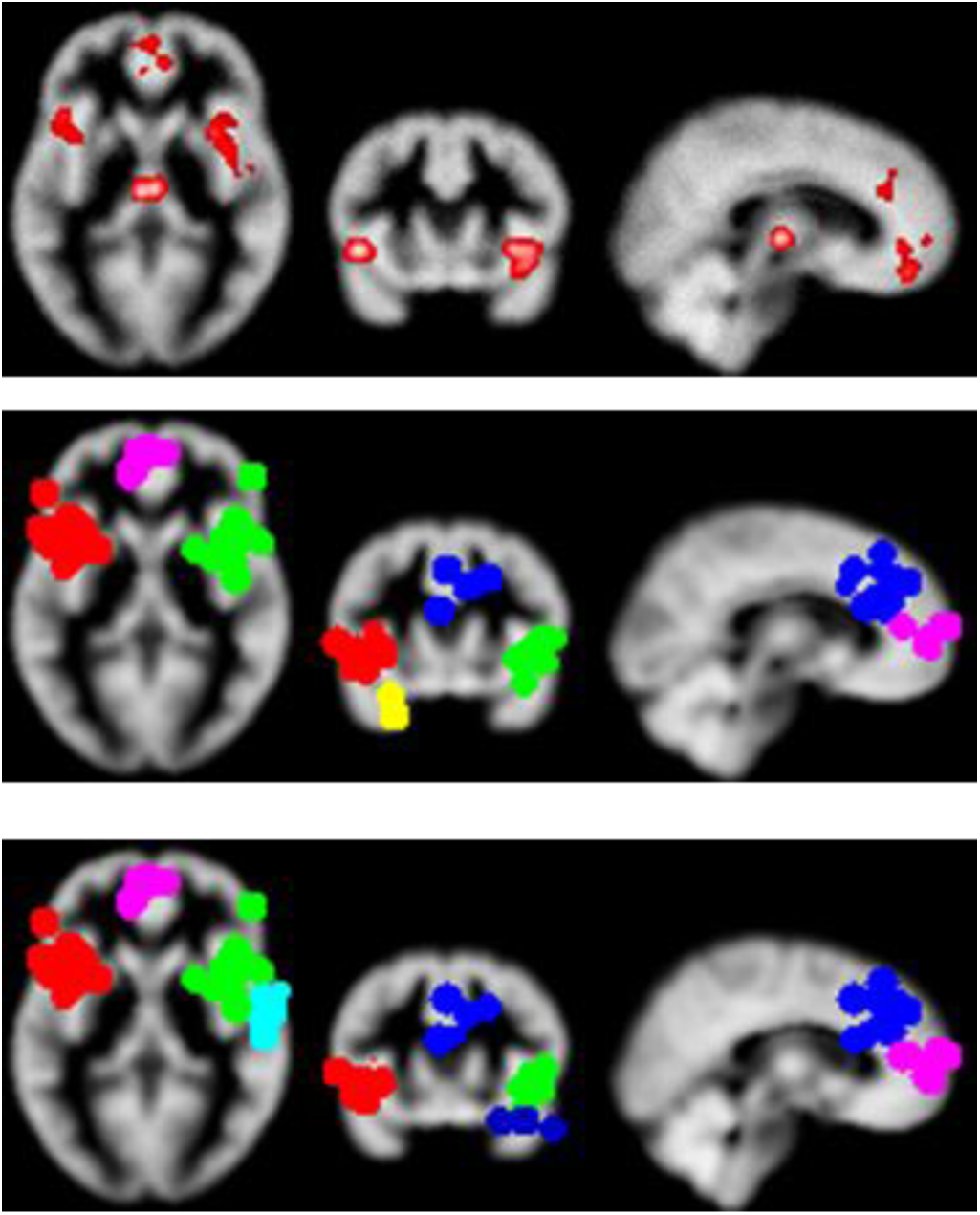
Results of the MA of VBM studies of Schizophrenia. ALE clusters (top), CBRES (middle) and CBMAN (bottom).

**Figure 5.**
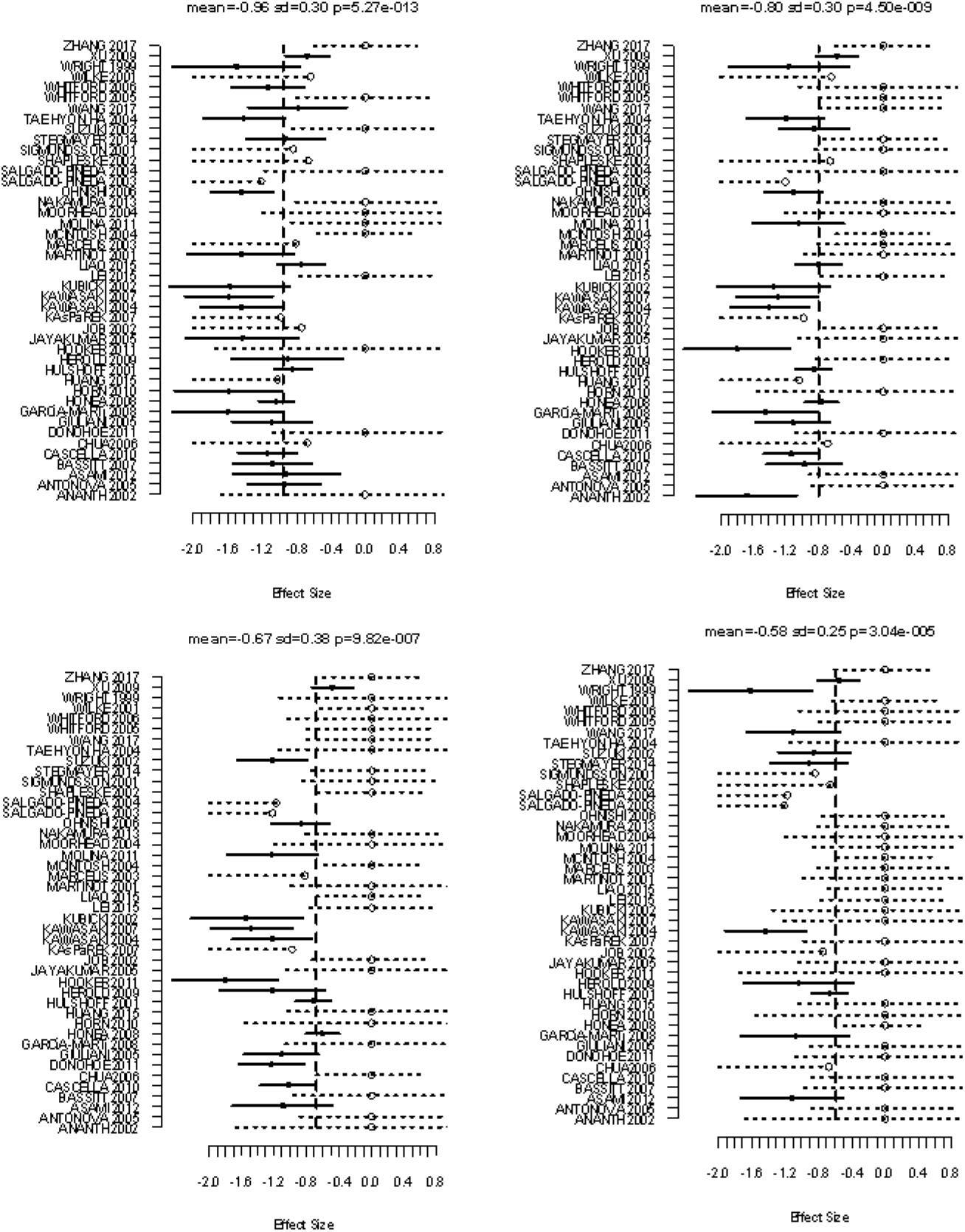
Forest plots of the 4 most significant clusters from the CBRES analysis. Solid lines indicate study specific sampling error of the standardised effect sizes. Dashed lines represent possible ranges of censored results.

**Figure 6.**
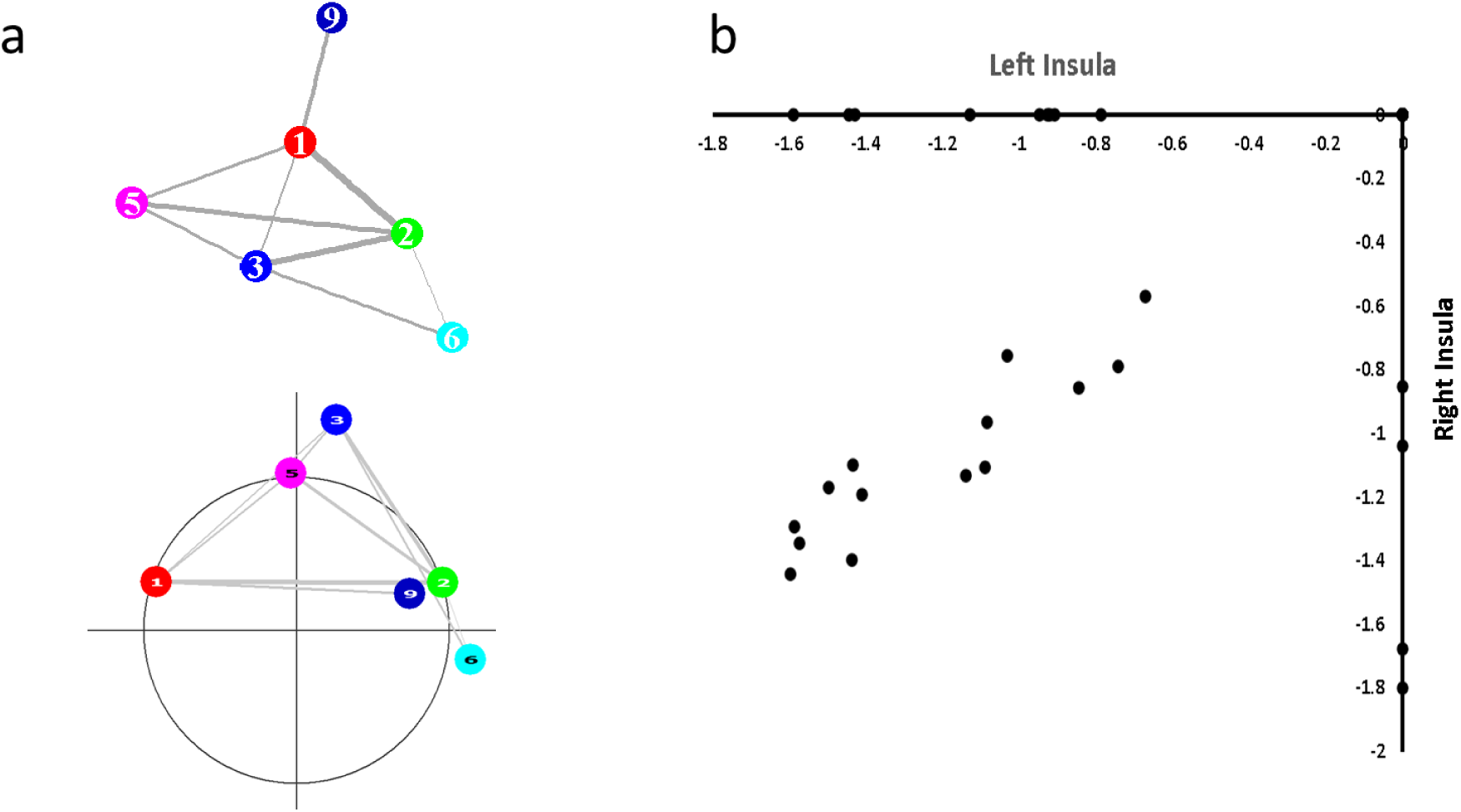
a) Clusters with significant connections according to CBMAN. Top shows a graph representation produced using R (R Development Core Team, 2008) and the iGraph package (Csardi and Nepusz, 2006), while bottom shows clusters placed such that the *x* and *y* locations are the Talairach *x* and *y* locations scaled such that the radius depends on the *z* Talairach coordinate (*z*=0 is indicated by the circle), which is generated by CBMAN. Covariance of effect sizes in clusters is represented by the thickness of the connecting lines. b) shows the scatter plot of the standardized effect sizes in the left and right insula. Circle markers on the x & y axes are censored.

For CBMAN analysis a minimum of 5 studies contributing uncensored results to a cluster is used as a constraint for statistical analysis because it is not possible to derive a p-value substantially less than 0.05 using the permutation test with fewer. To maintain comparability a constraint of 5 as a minimum number of studies in a cluster is also used in CBRES.

### Minimum density for valid cluster

To estimate the minimum density clusters of *n* coordinates are simulated, placed spatially at random with a spread drawn randomly from the empirical truncated *N(4,0.4^2^)* distribution. For each simulation the peak density (equation (1)) is recorded. The minimum density is the 5^th^ percentile of these recordings after 1000 simulations.

### Inclusion of region of interest studies

Some VBM or fMRI studies do not include a whole brain analysis, instead considering only a-priori hypothesised regions of interest. Preselection of these regions suggests there may be good evidence that they are relevant to the hypothesis, and ideally should be included in any meta-analysis. For CBMA methods that assume a null hypothesis of random coordinates, analysis of ROI studies violates this assumption (Müller et al., 2018). With methods that fit random effect models to the reported Z scores to estimate the mean effect (Albajes-Eizagirre et al., 2018; Costafreda, 2012; Tench et al., 2017), the assumption is that any unreported effects are censored by the study whole brain significance threshold. In ROI studies there is no such threshold, either the ROI analysis was conducted or it was not, so no assumption about censoring from a random effects model can be made. CBMAN does not use a null of random coordinates and the contribution of studies to clusters, rather than voxels, can be a special case where an ROI study not reporting is not considered at all rather than being considered censored. Consequently, CBMAN can analyse ROI studies, and this has been incorporated into the updated version.

### CBMA of Schizophrenia studies

A CBMA of Schizophrenia has previously been performed (Ananth et al., 2002) using the popular activation likelihood estimate (ALE) algorithm (Eickhoff et al., 2012; Laird et al., 2005; Turkeltaub et al., 2002). Here a similar analysis is performed, updated with more recent VBM studies of Schizophrenia. This should not be considered a meta-analysis of Schizophrenia, since the rigorous methods used in MA have not been strictly adhered to. The results are to demonstrate the output of CBRES and CBMAN with the new clustering algorithm. The latest version of GingerALE will also be used to produce the ALE results on the same data for comparison. CBMAN and CBRES are performed using an FDR method of type 1 error control. ALE is performed using cluster level FWE (0.05) with a p-value threshold of 0.001 and 1000 iterations. For comparison CBRES is also run with 1000 iterations rather than its default 4000.

Coordinates and study details are available at http://doi.org/10.17639/nott.7039.

## Results

### Minimum density for valid cluster

One thousand simulations were performed for each different number of coordinates forming the cluster. The minimum density was the 5^th^ percentile of the 1000 simulations. As a function of the number of coordinates the minimum density, *mD*, is

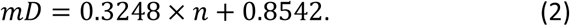

### Coordinate based meta-analysis of VBM studies of Schizophrenia

A total of 44 VBM studies of schizophrenia were included with a total of 514 coordinates. The studies involved a total of 1971 healthy controls and 1941 subjects with schizophrenia. The study density, computed using equation (1), and the clusters produced using the proposed clustering algorithm are shown in figure (3). One notable point is that the Thalamus is formed by two clusters, split left and right, despite the proximity. This resulted from the presence of two density peaks. The results of analysis by the ALE algorithm, CBRES, and CBMAN are depicted in figure (4). The cluster coordinates are also given in table (1).

Comparing the three analyses there is good agreement. The ALE and CBRES analyses are most similar and both test a null hypothesis of random coordinates. CBMAN misses a significant CBRES/ALE cluster (Parahippocampal gyrus) but identifies two extra significant clusters (Inferior frontal and Transverse Temporal gyrus) not detected by CBRES or ALE. These differences probably reflect the different null hypothesis tested by CBMAN. Another difference is the significant thalamic cluster detected by the ALE algorithm but not by CBRES or CBMAN. This might be because the voxel-wise ALE algorithm does not split the thalamic cluster into left and right clusters, despite reporting separate left and right ALE peaks. Merging the bilateral thalamic clusters into one can increase the number of studies contributing and consequently the statistical significance. It may be that the significant thalamic cluster is a consequence only of this merging, with no statistical evidence for individual left and right thalamic clusters. Indeed, the two thalamic clusters are only significant in CBRES with FCDR>0.3, and the connection between the two according to CBMAN has a p-value of 0.54.

There is one further consideration of these results. The left Parahippocampal Gyrus is declared significant by both ALE and CBRES. However, it is only declared significant using 1000 coordinate randomisations to define the null hypothesis. In CBRES 1000 randomisations is inadequate to fully describe the null hypothesis and at 4000 (the default) it is no longer significant. While CBRES takes only a few hours to execute, ALE would have taken 8 days to execute 4000 iterations as it needs to compute its test statistic for every voxel rather than every cluster. Since both methods use coordinate randomisations to define the null it is likely that the default 1000 randomisations used by ALE is inadequate; note that while 1000 iterations was considered convergent for ALE (Eickhoff et al., 2012), an implementation issue (Eickhoff et al., 2016a) was subsequently discovered that calls this into question.

### Further CBRES and CBMAN analysis

While CBMA can produce voxel-wise maps of significant spatial consistency of reported coordinates across studies testing the same hypothesis, using a cluster-wise analysis makes possible further analysis. Figure (5) shows forest plots of the 4 most significant clusters declared significant by CBRES. These are useful to inspect the heterogeneity of the reported effects and to diagnose any issues with the data; it is particularly easy to make errors when record the tabulated data from many independent studies. For CBMAN a map of network like connections, and the standardised Z score effect size correlation inspected (figure (6)). Network analysis has also been used on such results (Cauda et al., 2018; Lancaster et al., 2005; Neumann et al., 2005; Tatu et al., 2018), offering alternatives to CBMA.

## Discussion

Neuroimaging studies often involve complex analysis with methods that are still evolving. They can be limited by small samples and inconsistent application of methodology, resulting in often difficult to interpret results. Coordinate based meta-analysis offers the chance to summarise the results from multiple independent neuroimaging studies that have tested a common hypothesis, with the results indicating effects that are in some way statistically consistent. CBMA also has limitations, two of which are that the voxel-wise analysis necessitates the use of conservative FWE type 1 error control and prevents the inclusion of region of interest studies. These can be overcome using cluster-wise analysis. Here a clustering algorithm based on DENCLUE has been detailed.

The aim of the new clustering algorithm is to form a smooth function consisting of high, single peak, density regions. Initialising the recursive clustering algorithm at each reported coordinate and traversing the smooth function in the direction of increasing density finds these peaks and assigns coordinates to them to form clusters. Once study membership to each cluster is established, several analysis approaches not available with voxel-wise analysis become available. False discovery rate can be used as a less conservative alternative to FWE (the best option for voxel-wise analysis) and has the benefit that the error rate is estimated in terms of the proportion of significant results (clusters). Cluster-wise analysis also makes it possible to use cooccurrence analysis, with a hypothesis that the multiple reported peak coordinates form a consistent pattern of activation or grey matter alteration by testing a null hypothesis of no cooccurrence between different brain regions; there is also a voxel-gwise cooccurrence algorithm but it tests a different hypothesis of spatially random coordinates (Chu et al., 2015). Some algorithms achieve this using just coordinates (no Z scores) by forming clusters from reported coordinates within a fixed spherical region placed at locations identified using the ALE algorithm (Cauda et al., 2018; Tatu et al., 2018), or within ALE clusters themselves (Lancaster et al., 2005; Neumann et al., 2005). There are problems with this approach in that: cooccurrence and CBMA analyses can involve different regions since they test different hypotheses (see table 1) so some regions might be missed, and selecting nodes based on the highest ALE preselects the nodes of highest occurrence, which is not independent of cooccurrence and might be considered ‘double-dipping’ (Kriegeskorte et al., 2009). In contrast CBMAN measures cooccurrence using correlated Z scores between all valid clusters, rather than the coincidence of simultaneous reporting by studies in multiple clusters, side-stepping these problems. Finally, cluster-wise cooccurrence methods can incorporate ROI studies thus increasing the pool of available data to analyse. This is important because ROI studies may have used strong a-priori evidence to select the regions making the results particularly relevant to the MA. The new clustering method reported here has been incorporated into CBRES and CBMAN methods such that the results from each are directly comparable.

**Table 1.** Peak coordinates of clusters from CBMAN, CBRES, and ALE analyses on the Schizophrenia coordinate data. A dash (-) indicates no significant cluster. Coordinate system: Talairach. Note that the CBRES cluster marked by a * is only found for 1000 coordinate randomisations as used by ALE. Using the default 4000 randomisations produces convergent results and eliminates this cluster.

| Location | CBMAN | CBRES | ALE |
| --- | --- | --- | --- |
| L. Insula. BA 13 | -40.0, 14.0, -2.0 | -40.0, 14.0, -2.0 | -40.1,8.1,-1.4 |
| R. Insula. BA 13 | 42.0, 14.0, 0.0 | 42.0, 14.0, 0.0 | 41,13.6,-1.4 |
| R. Cingulate gyrus. BA 32 | 6.0, 32.0, 26.0 | 6.0, 32.0, 26.0 | 5.2,32.4,27.3 |
| R. Inferior frontal gyrus. BA 47 | 30.0, 10.0, -16.0 | - | - |
| R. Transverse Temporal Gyrus | 60.0, -10.0, 10.0 | - | - |
| L. Medial Frontal Gyrus. BA 11 | -2.0, 50.0, 2.0 | -2.0, 50.0, 2.0 | -1.6,44.2,-4.3 |
| L. Parahippocampal Gyrus. BA 34 | - | -18.0, -7.0, -19.0* | -19.2,-5.1,-19.2 |
| Thalamus | - | - | -1.4,-17,5.5 |

## Summary

A clustering algorithm based on the empirical observation that the spread of coordinates within typical coordinate-based analyses is around 4mm has been presented. Using this the studies contributing to each cluster can be established. This has been incorporated into updated CBRES and CBMAN algorithms available in NeuRoi software. Using a cluster-wise rather than a voxel-wise schemes have several advantages: network like cooccurrence analysis, FDR rather than FWE type 1 error control, and the option to include important region of interest studies.

## Appendix Assigning coordinates to local study density peaks

~~~
Global variables: **PeakVoxel, PeakDensity**
Initialise **PeakDensity** to 0
Define Image **Visited**[] and initialise all voxels to 0//*Prevent repeated calculation*
Function: **FindPeakDensity**(start_voxel, seed_density)
{
    D=**Density**(start_voxel); //*use Equation (1)*
    **Visited**[start_voxel]=1;
    If (D>**PeakDensity**) {
        **PeakDensity**=D; //*D is highest so far so is proposed peak*
        **PeakVoxel**=start_voxel;
    }
If (D>seed_density){
        For all neighbours n of start voxel:
            If (**Visited**[n]==0) **FindPeakDensity**(n, D);//*Continue looking for peak*
    }
}
Output: The density peak is at **PeakVoxel** and has a density=**PeakDensity**.
~~~

Cluster finding algorithm

- For each coordinate call **FindPeakDensity** and assign the coordinate to a **PeakVoxel**
- Each of the identified **PeakVoxel** and the assigned coordinates are proposed clusters
- For each proposed cluster remove all but the closest contribution from each study
- If a proposed cluster has more than the minimum number of studies contributing, and the **PeakDensity** is greater than the minimum density (equation (2) for 4 coordinates), the cluster is valid.

